# Type I interferon signaling promotes early innate control of *Borrelia burgdorferi* infection

**DOI:** 10.64898/2026.02.13.705697

**Authors:** Raj Priya, Sajith Raghunandanan, Darco Mihaljica, Nada S. Alakhras, Mark H. Kaplan, X. Frank Yang

## Abstract

*Borrelia burgdorferi*, the causative agent of Lyme disease, elicits a robust type I interferon (IFN-I) response that has been strongly associated with chronic inflammatory manifestations such as Lyme arthritis. Although IFN-I is induced early after infection, its functional contribution during the initial stages of *B. burgdorferi* infection has remained unclear. Here, we identify a critical protective role for IFN-I signaling in the early control of *B. burgdorferi*. At 5 days post-infection, mice lacking the IFN-I interferon receptor (IFNAR1) exhibited markedly elevated spirochetal burdens at the site of inoculation, indicating that IFN-I is required to restrict early bacterial expansion. By 10 days post-infection, IFNAR1 deficiency resulted in significantly increased bacterial loads in skin and heart tissues, while joint burdens remained unaffected. Mechanistically, IFN-I signaling enhanced macrophage phagocytosis and intracellular killing of *B. burgdorferi* both *in vitro* and *in vivo*, and promoted pro-inflammatory cytokine production and recruitment of innate immune cells to the site of infection. Conversely, pharmacological activation of IFN-I signaling augmented macrophage antimicrobial function and improved bacterial clearance. Together, these findings establish IFN-I as an essential component of early host defense against *B. burgdorferi* and reveal a time-dependent role for IFN-I signaling in Lyme disease pathogenesis, with protective functions during early infection that contrast with its pathogenic effects at later stages.

**AUTHOR SUMMARY:** Lyme disease is caused by the spirochete *Borrelia burgdorferi*, which is transmitted to humans through the bite of infected ticks. Soon after infection, the spirochete must evade the host’s early immune defenses in the skin to disseminate to other tissues and cause disease. Type I interferons (IFN-I) are best known for their antiviral roles, but they are also strongly induced during Lyme disease. While IFN-I responses have been linked to the development of Lyme Arthritis in later stages of infection, their role during the earliest phase of Lyme disease has remained unclear. In this study, we show that IFN-I interferon signaling plays a critical protective role early after *B. burgdorferi* infection. Mice lacking the IFN-I interferon receptor were unable to efficiently control bacterial growth at the site of infection and showed increased bacterial dissemination. Mechanistically, IFN-I signaling enhanced the ability of macrophages to engulf and kill *B. burgdorferi*, promoted inflammatory cytokine production, and supported the recruitment of immune cells to the site of infection. Importantly, short-term activation of IFN-I signaling enhanced bacterial clearance, whereas loss of this pathway impaired early immune control. Together, these findings reveal that IFN-I has time-dependent roles during Lyme disease, protecting the host during early infection while contributing to inflammation at later stages. By defining the time-dependent role of IFN-I signaling in Lyme disease, our study advances understanding of how innate immune responses shape infection outcome and highlights potential strategies for enhancing early host defense.

## INTRODUCTION

Lyme disease, caused by the spirochete *Borrelia burgdorferi (B. burgdorferi)*, is the most prevalent vector-borne illness in the United States and Europe [1]. Each year, hundreds of thousands of cases are reported, underscoring its substantial public health burden [2, 3]. After transmission, *B. burgdorferi* establishes infection in the skin, where it evades early immune clearance and disseminates to distant tissues, including the joints, heart, and nervous system [4, 5]. Disseminated infection can lead to severe clinical outcomes such as Lyme arthritis, Lyme carditis, and neuroborreliosis, with Lyme arthritis being the most common late-stage manifestation. The absence of a human vaccine, combined with the challenges in diagnosing and managing persistent disease, underscores the critical need to understand the host-pathogen interactions that shape disease outcome. A crucial gap exists in understanding the initial innate immune response at the site of infection, where the balance between bacterial clearance and early dissemination is established. *B. burgdorferi’s* ability to survive and disseminate relies on successfully overcoming the host’s initial defenses. Thus, defining early immune responses at the site of infection is essential for understanding the mechanisms that determine bacterial clearance versus persistence and chronic inflammation.

IFN-I, classically defined by their roles in antiviral defense [6], are rapidly induced upon nucleic acid sensing and activate hundreds of interferon-stimulated genes (ISGs) that establish broad antimicrobial states [7, 8]. IFN-I is now recognized to modulate immune responses during bacterial infections as well [9]. However, IFN-I responses exert complex and sometimes opposing effects during bacterial infections, with outcomes that vary depending on the nature of a pathogen, infection site, timing, and strength of induction. In some contexts, IFN-I supports antibacterial immunity by enhancing immune cell activation and increasing inflammatory cytokines that contribute to bacterial control [10–13]. However, IFN-I can also be detrimental to the host, by suppressing protective immune responses through the induction of anti-inflammatory cytokines and inhibition of proinflammatory mediators and chemokines, or promoting apoptosis of macrophages and lymphocytes, resulting in increased bacterial dissemination [14–16].

*B. burgdorferi*, an extracellular spirochetal pathogen, elicits a robust IFN-I response in both mice and humans [17, 18]. Much of this IFN-I induction occurs through pathways independent of MyD88- and TRIF-mediated Toll-like receptors (TLRs) [18–21], but is driven largely by intracellular sensing of spirochetal components through the cGAS–STING axis [22, 23]. In addition, endosomal TLR7/8 sensing of *B. burgdorferi* RNA may also contribute to IFN-I production in certain host cells [17, 24, 25]. *B. burgdorferi*-induced IFN-I plays a pathogenic role in Lyme disease. Research to date has focused primarily on its effects during the later stages of infection, where elevated IFN-I levels correlate with increased inflammation and tissue damage, which contributes significantly to the development and severity of Lyme arthritis [21, 26–29]. In addition, IFN-I responses drive the abnormal accumulation of naïve B cells in lymph nodes, which compromises the efficacy of the adaptive immune response [20].

Accumulated evidence indicates that during the early acute stage of infection, *B. burgdorferi* also induces a strong IFN-I response in mouse models, tick bite sites, and human erythema migrans lesions [30–33]. Transcriptional profiling of human erythema migrans (EM) lesions reveals that ISGs constitute a dominant portion of the inflammatory signature, preceding the onset of adaptive immunity [32]. Increased IFN-I in serum and EM blister fluids correlates with disseminated disease in patients, and *B. burgdorferi* phagocytosis induces IFN-I production from macrophages, plasmacytoid dendritic cells (pDCs), and myeloid dendritic cells (mDCs) [24, 31, 32]. In murine infection models, *B. burgdorferi* triggers the expression of IFN-β and multiple ISGs in skin and blood within the first days of infection, and the magnitude of this early IFN response correlates with strain dissemination capacity [31]. Despite these observations, the role of IFN-I responses during the early stage of *B. burgdorferi* infection remains unclear.

To address this critical unresolved question, in this study, we investigated the role of IFN-I signaling in the early control of *B. burgdorferi* infection. Specifically, we evaluated its influence on phagocytosis, intracellular killing, production of proinflammatory cytokines, immune cell recruitment, and bacterial burden at the site of infection. We demonstrate that IFN-I is crucial for effective early control of *B. burgdorferi* by enhancing bacterial uptake and augmenting macrophage-derived antimicrobial responses, thereby significantly reducing the bacterial burden during the initial phase of infection. These results reveal a previously underappreciated protective mechanism driven by IFN-I that coordinates effective early innate immunity. By identifying IFN-I as a central regulator of early phagocytic and antimicrobial responses, this work provides insights suggesting that therapeutic strategies designed to enhance IFN-I signaling during the initial stages of infection could strengthen host defense, limit bacterial burden, and potentially reduce the risk of post-treatment complications.

## RESULTS

### IFN-I signaling controls early *B*. *burgdorferi* burden

To determine the role of IFN-I signaling during early *B. burgdorferi* infection, we quantified spirochetal load in skin, heart, and joint tissues of wild-type (WT) and *Ifnar1*⁻/⁻ mice at 5- and 10-days post-infection (DPI) using quantitative PCR (qPCR) **(Fig. 1**). At the site of inoculation, *Ifnar1*⁻/⁻ mice exhibited a significantly higher *B. burgdorferi* burden compared with WT controls, with 3.2-fold and 24-fold increases at 5 and 10 DPI, respectively (**Fig. 1A**, left). In addition, spirochetal burdens were assessed at distal sites, including the heart and joints. At 10 DPI, there was a significant increase in spirochetal burden in the heart tissues of *Ifnar*1⁻/⁻ mice (**Fig. 1B**, center). In joint tissues, spirochetal burdens were comparable between WT and *Ifnar*1⁻/⁻ mice (**Fig. 1A** and **B**, right). These results demonstrate that IFN-I signaling contributes to the early control of *B. burgdorferi* replication by limiting bacterial burden at the site of infection and in the distal heart tissues.

**Fig. 1.**
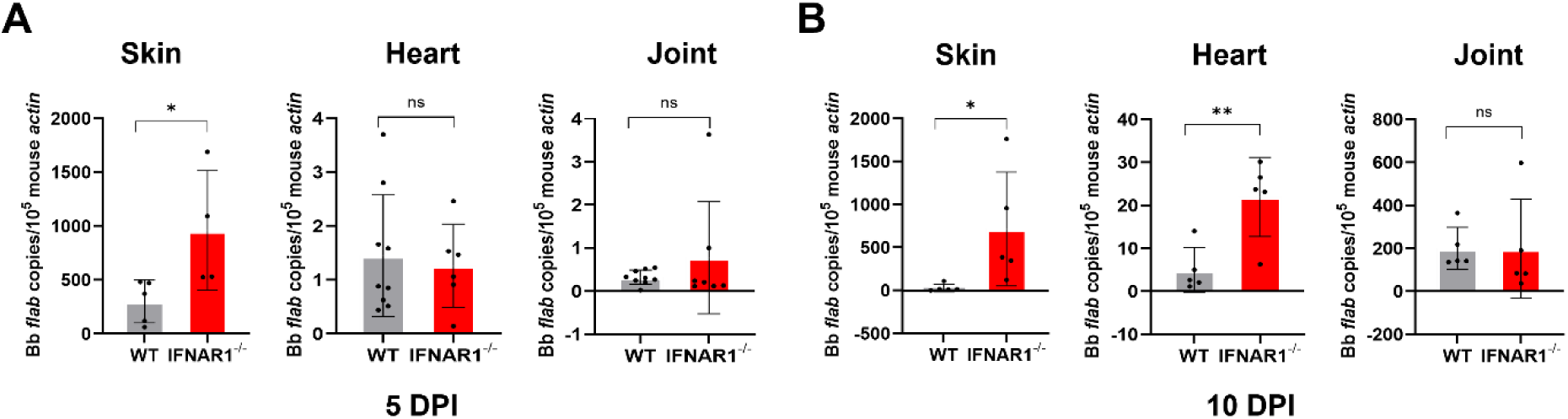
IFN-I signaling contributes to early control of *B. burgdorferi* burden. Wild-type (WT) and *Ifnar*1⁻/⁻ mice were intradermally infected with *B. burgdorferi* (1 × 10^4^ spirochetes/mouse). At 5- and 10-DPI, skin (site of infection), heart, and joint tissues were harvested, DNA was isolated, and *B. burgdorferi* burden was quantified by quantitative PCR (qPCR). The number of *B. burgdorferi flaB* copies was normalized to mouse *β-actin*. Data are presented as mean ± SD (n = 4–10 mice/ group). Statistical significance was determined using a two-tailed unpaired Student’s *t*-test. *, *P* < 0.05; **, *P* < 0.01.

### IFN-I signaling enhances macrophage phagocytosis of *B. burgdorferi in vitro*

To investigate the mechanisms by which IFN-I signaling contributes to early control of *B. burgdorferi*, we first examined macrophage phagocytosis *in vitro*. Bone marrow–derived macrophages (BMDMs) from WT and *Ifnar1*⁻/⁻ mice, as well as RAW 264.7 macrophages treated with or without an IFNAR1-blocking antibody, were infected with GFP-expressing *B. burgdorferi* at a multiplicity of infection (MOI) of 10 for 2.5 hours. Flow cytometric analysis revealed a significant reduction in bacterial uptake by *Ifnar1*⁻/⁻ BMDMs compared with WT controls, as indicated by both a decreased percentage of GFP-positive cells and reduced GFP mean fluorescence intensity (MFI) (**Fig. 2A**). Similarly, blockade of IFNAR1 in RAW 264.7 macrophages resulted in a marked decrease in GFP-positive cells and GFP MFI relative to isotype-treated controls (**Fig. 2B**). Confocal microscopy confirmed these findings, showing reduced intracellular GFP signal in *Ifnar1*⁻/⁻ BMDMs and in RAW 264.7 cells treated with IFNAR1-blocking antibody (**Fig. 2C**). Together, these results demonstrate that IFN-I signaling is required for efficient macrophage-mediated phagocytosis of *B. burgdorferi in vitro*.

**Fig. 2.**
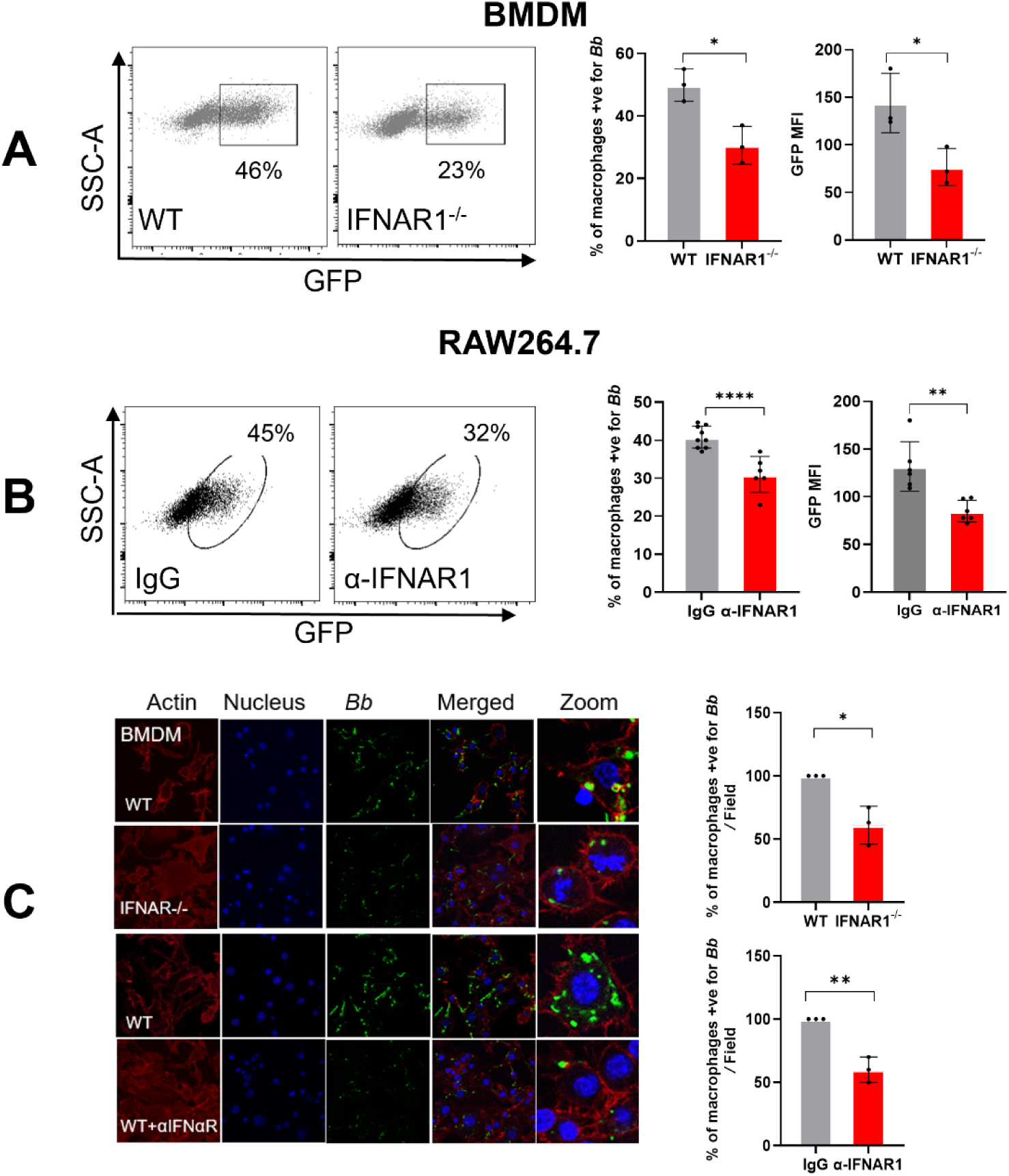
**IFN-I signaling enhances macrophage-mediated phagocytosis of *B. burgdorferi in vitro***. Bone marrow–derived macrophages (BMDMs) from wild-type (WT) and *Ifnar1*⁻/⁻ mice, as well as RAW 264.7 macrophages treated with or without an IFNAR1-blocking antibody, were infected with GFP-expressing *B. burgdorferi* at an MOI of 10 for 2.5 hours. (**A, B**) Flow cytometric analysis of phagocytosis. Representative flow cytometry plots (SSC versus GFP) and quantification of the percentage of GFP-positive cells and GFP MFI in (**A**) BMDMs and (**B**) RAW 264.7 cells. (**C**) Confocal microscopy analysis. Representative confocal images showing GFP-*B. burgdorferi* (green) within BMDMs and RAW 264.7 macrophages, with quantification of GFP-positive cells. Data are presented as mean ± SD from at least three independent experiments. Statistical significance was determined using a two-tailed unpaired Student’s *t*-test. *, *P* < 0.05; **, *P* < 0.01; ****, *p* < 0.0001.

### IFN-I signaling promotes macrophage phagocytosis of *B. burgdorferi in vivo*

To determine whether IFN-I signaling contributes to macrophage phagocytosis of *B. burgdorferi in vivo*, WT and *Ifnar1*⁻/⁻ mice were infected intraperitoneally with GFP-expressing *B. burgdorferi*. Peritoneal exudate cells were harvested 6 hours post-infection (HPI), and macrophage phagocytosis was assessed by the presence of GFP signal in macrophages by flow cytometry as described previously [34] **(Fig. 3A)**. Macrophages were identified as live CD45⁺CD11b⁺F4/80⁺ cells using a sequential gating strategy (**Fig. 3B**). Representative flow cytometry plots demonstrated reduced GFP signal (GFP versus SSC) in peritoneal macrophages from *Ifnar1*⁻/⁻ mice compared with WT controls (**Fig. 3C**). Quantitative analysis revealed a significantly lower percentage of GFP-positive macrophages in *Ifnar1*⁻/⁻ mice relative to WT mice (**Fig. 3D**), indicating impaired bacterial uptake in the absence of IFN-I signaling.

**Fig. 3.**
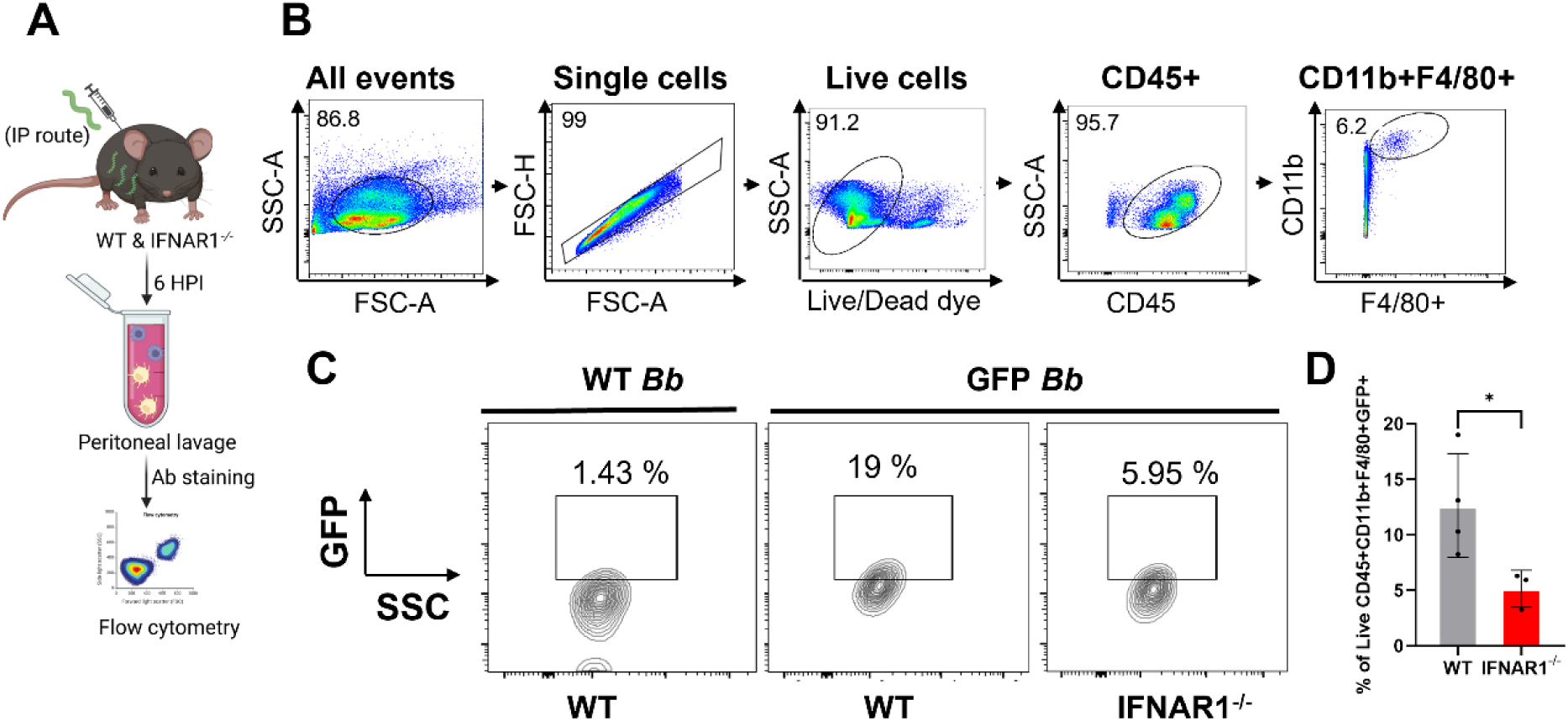
**IFN-I signaling promotes macrophage-mediated phagocytosis of *B. burgdorferi in vivo***. Wild-type (WT) and *Ifnar1*⁻/⁻ mice were infected intraperitoneally with GFP-expressing *B. burgdorferi* (1 × 10⁷ spirochetes/ mouse), or with non-fluorescent *B. burgdorferi* (as a control for autofluorescence). Peritoneal exudate cells (PECs) were harvested at 6 HPI and stained with a viability dye, anti-CD45, anti-CD11b, and anti-F4/80 antibodies. Phagocytosis was assessed by flow cytometry to detect the GFP signal in live CD45^+^CD11b^+^F4/80^+^ peritoneum macrophages. (**A**) Experimental workflow illustrating intraperitoneal infection, peritoneal cell isolation, and flow cytometric analysis. (**B**) Flow cytometry gating strategy used to identify live peritoneal macrophages. (**C**) Representative flow cytometry plots (GFP vs SSC) showing GFP signal in peritoneal macrophages from WT and *Ifnar1*⁻/⁻ mice. (**D**) Quantification of the percentage of GFP-positive peritoneal macrophages. Data are presented as mean ± SD (n = 3-4 mice/group). Statistical significance was determined using a two-tailed unpaired Student’s *t-*test. *, *P* < 0.05.

#### IFN-I signaling promotes intracellular killing and lysosomal targeting of *B. burgdorferi*

To determine whether IFN-I signaling influences macrophage antimicrobial activity beyond bacterial uptake, we next assessed intracellular killing of *B. burgdorferi* using a gentamicin protection assay. BMDMs from WT and *Ifnar1*⁻/⁻ mice were infected with *B. burgdorferi*, extracellular bacteria were eliminated by gentamicin treatment, and intracellular bacterial viability was quantified by limiting dilution in 96-well plates. Lysates from *Ifnar1*⁻/⁻ BMDMs yielded a significantly higher proportion of *B. burgdorferi*–positive wells compared with WT macrophages, indicating impaired intracellular killing in the absence of IFN-I signaling (**Fig. 4A**). To further examine the mechanism underlying this defect, we evaluated lysosomal targeting of internalized spirochetes by assessing colocalization with the lysosomal marker LAMP1 using confocal microscopy. *Ifnar1*⁻/⁻ macrophages exhibited a reduced level of LAMP1 colocalization with internalized *B. burgdorferi* relative to WT cells (**Fig. 4B**). Together, these findings indicate that IFN-I signaling promotes efficient intracellular killing of *B. burgdorferi* by facilitating lysosomal targeting following phagocytosis.

**Fig. 4.**
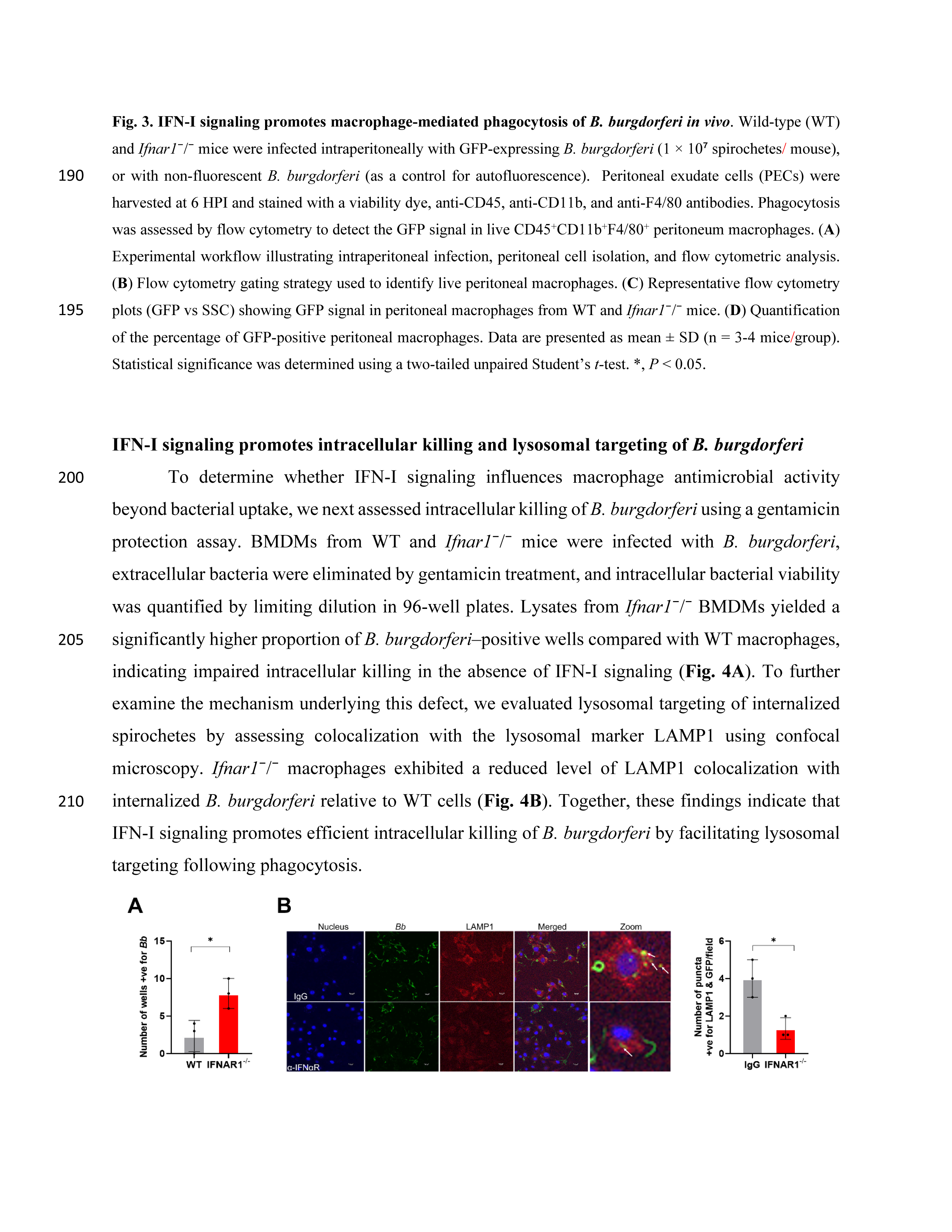
IFN-I signaling promotes the intracellular killing and lysosomal targeting of *B. burgdorferi*. **(A)** Gentamicin protection assay. Bone marrow–derived macrophages (BMDMs) from wild-type (WT) and *Ifnar1*⁻/⁻ mice were infected with *B. burgdorferi* for 2.5 hours, followed by treatment with gentamicin (200 µg/mL) to eliminate extracellular bacteria. Following washing and lysis, cell lysates were diluted in BSK-II medium and plated in 96-well plates. Intracellular bacterial viability was assessed by limiting dilution, and data are presented as the number of *B. burgdorferi*-positive wells. (**B**) Lysosomal targeting of internalized *B. burgdorferi*. Colocalization of internalized spirochetes with the lysosomal marker LAMP1 was assessed by confocal microscopy. Representative images show *B. burgdorferi* (**green**), LAMP1 (**red**), and colocalization (**yellow**). Quantification represents the percentage of internalized bacteria that colocalize with LAMP1. Data are presented as mean ± SD from three independent experiments. Statistical significance was determined using a two-tailed unpaired Student’s *t*-test. *, *P* < 0.05.

#### Exogenous IFN-β enhances macrophage phagocytosis and intracellular killing of *B. burgdorferi*

To determine whether activation of IFN-I signaling is sufficient to enhance macrophage antimicrobial function, we examined the effects of exogenous IFN-β on macrophage uptake and intracellular killing of *B. burgdorferi in vitro*. RAW 264.7 macrophages were infected with GFP-expressing *B. burgdorferi* in the presence or absence of recombinant IFN-β, and bacterial uptake was assessed by flow cytometry and confocal microscopy (**Fig. 5A**). Flow cytometric analysis demonstrated that IFN-β treatment significantly increased both the percentage of GFP-positive macrophages and the GFP MFI compared with untreated controls, indicating enhanced bacterial uptake (**Fig. 5B**). Consistent with these findings, confocal microscopy revealed increased intracellular GFP signal in IFN-β–treated macrophages, with a higher proportion of GFP-positive cells relative to untreated controls (**Fig. 5C**).

**Fig. 5.**
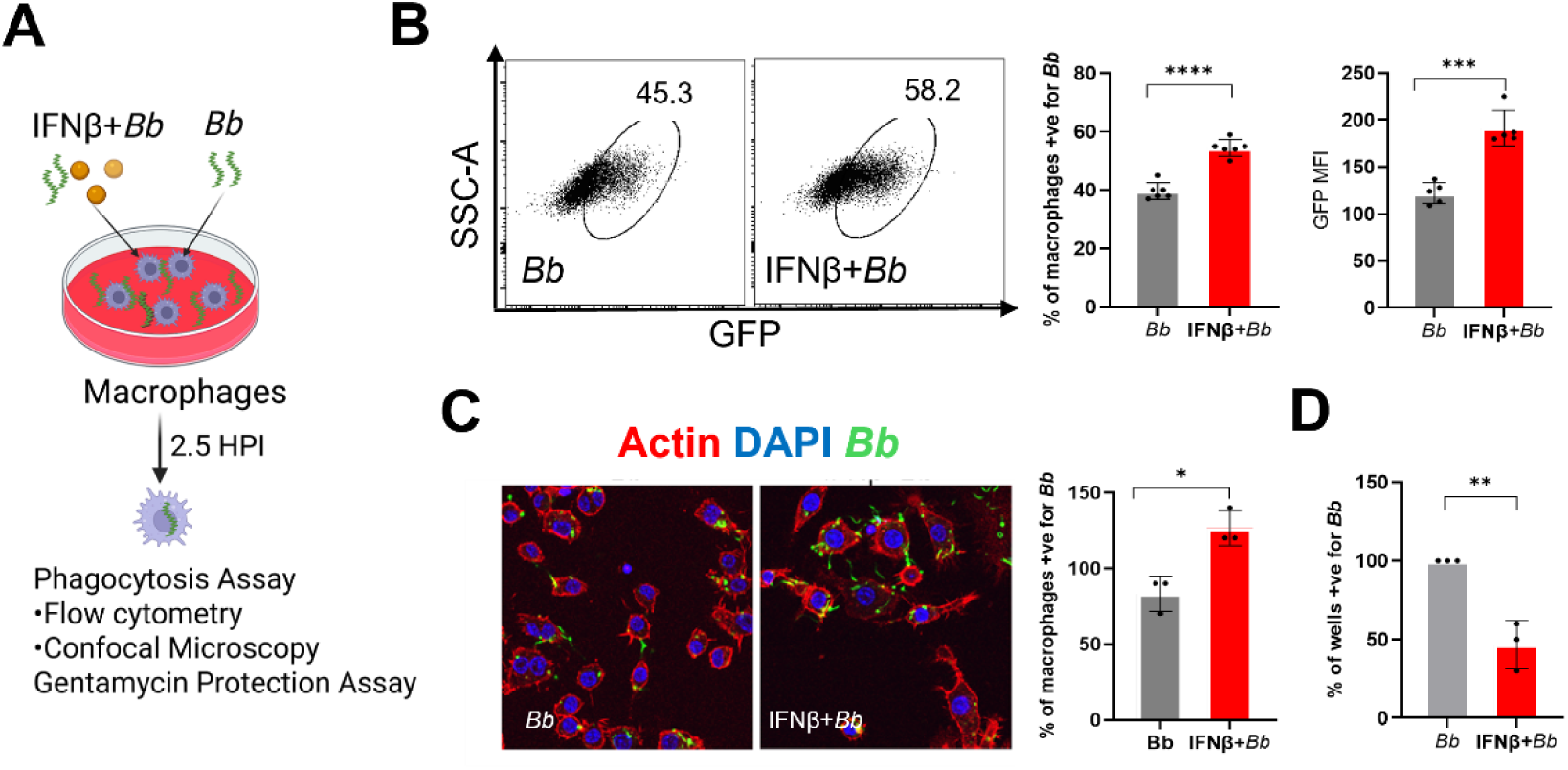
Exogenous IFN-β enhances macrophage phagocytosis and intracellular killing of *B. burgdorferi*. RAW 264.7 macrophages were infected with GFP-expressing *B. burgdorferi* at an MOI of 10 for 2.5 hours in the presence or absence of exogenous recombinant IFN-β (500 pg/mL). (**A**) Experimental schematic outlining IFN-β treatment, infection, and downstream analyses. (**B**) Flow cytometric analysis of phagocytosis. Representative plots and quantification of the percentage of GFP-positive macrophages and GFP mean fluorescence intensity (MFI). (**C**) Confocal microscopy analysis. Representative images showing intracellular GFP-*B.burgdorferi* and quantification of GFP-positive macrophages. (**D**) Gentamicin protection assay assessing intracellular bacterial viability following IFN-β treatment. Data are presented as mean ± SD from three independent experiments. Statistical significance was determined using a two-tailed unpaired Student’s *t-*test. *, *P* < 0.05; **, *P* < 0.01; ***, *P* < 0.001; ****, *P* < 0.0001.

To assess whether IFN-β treatment also affected intracellular bacterial viability, we performed a gentamicin protection assay following infection. IFN-β–treated macrophages exhibited a reduced frequency of culture-positive wells compared with untreated cells, indicating enhanced intracellular killing of *B. burgdorferi* (**Fig. 5D**). Together, these results demonstrate that activation of IFN-I signaling by exogenous IFN-β is sufficient to enhance macrophage phagocytosis and intracellular killing of *B. burgdorferi in vitro*.

#### IFN-I signaling modulates pro-inflammatory cytokine production in macrophages in response to *B. burgdorferi*

To determine whether impaired phagocytosis in the absence of IFN-I signaling alters subsequent downstream inflammatory responses, we measured pro-inflammatory cytokine production by macrophages following *B. burgdorferi* infection. BMDMs from WT and *Ifnar1*⁻/⁻ mice were infected with *B. burgdorferi*, and levels of IL-1β, IL-6, and TNF-α in culture supernatants were quantified at 3-, 6-, and 24-HPI. At 24 HPI, *Ifnar1*⁻/⁻ BMDMs exhibited a marked reduction in IL-1β and TNF-α production compared with WT macrophages (**Fig. 6A**). IL-6 production was also reduced in *Ifnar1*⁻/⁻ BMDMs at earlier time points, although levels approached those of WT cells by 24 HPI. These results indicate that IFN-I signaling contributes to optimal induction of key pro-inflammatory cytokines in murine macrophages in response to *B. burgdorferi*.

**Fig. 6.**
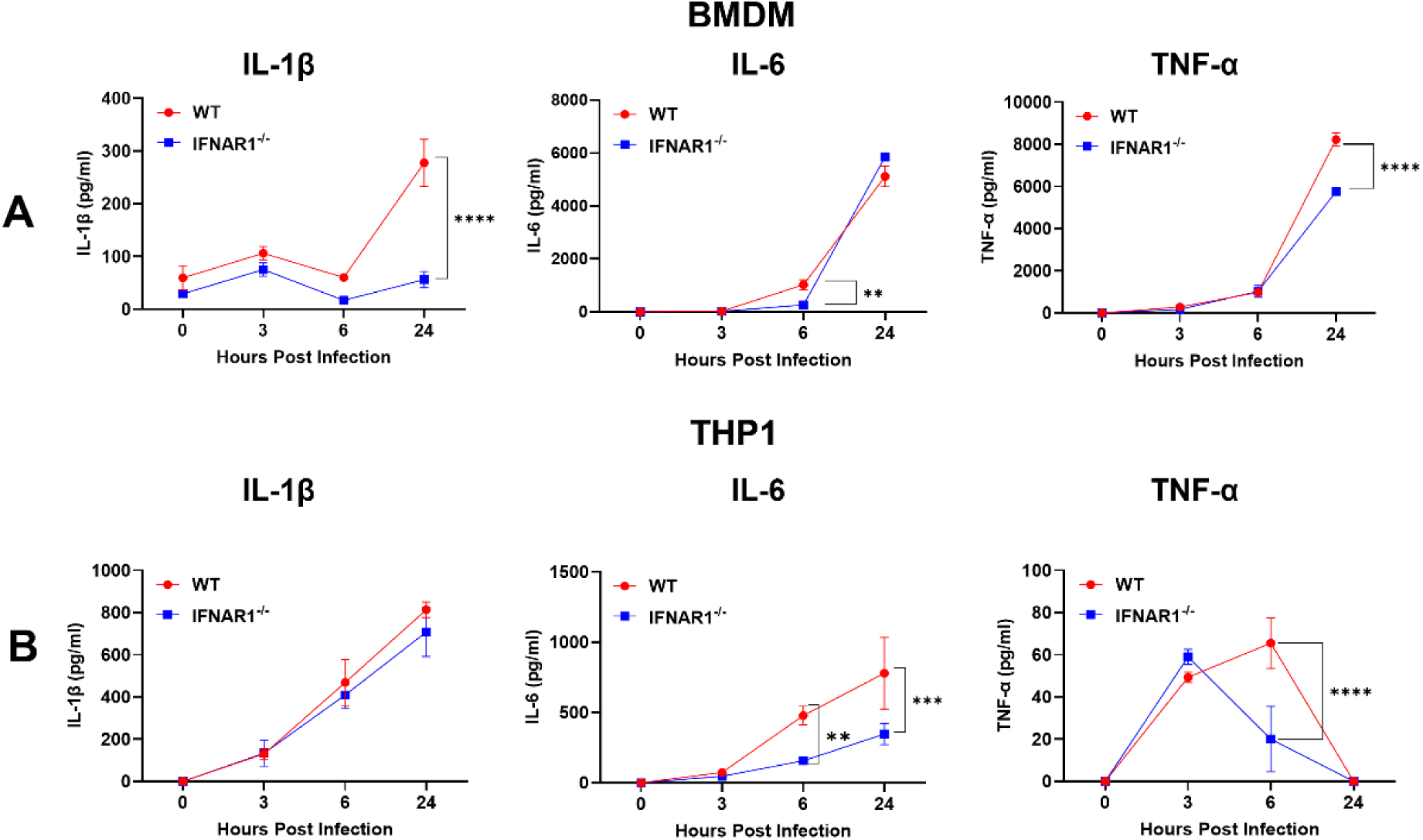
IFN-I signaling modulates pro-inflammatory cytokine production in macrophages following *B. burgdorferi* infection. (**A**) Murine macrophages. Bone marrow–derived macrophages (BMDMs) from wild-type (WT) and *Ifnar1*⁻/⁻ mice were infected with *B. burgdorferi* at an MOI of 10 for 3, 6, and 24 hours. Culture supernatants were collected, and levels of IL-1β, IL-6, and TNF-α were measured by ELISA. (**B**) Human macrophages. THP-1–derived macrophages were infected with *B. burgdorferi* in the presence or absence of an IFNAR1-blocking antibody. Cytokine levels in culture supernatants were quantified by ELISA at the indicated time points. Data are presented as mean ± SD from three independent experiments. Statistical significance was determined using a two-tailed unpaired Student’s *t-*test. **, *P* < 0.01; ***, *P* < 0.001; ****, *P* < 0.0001.

To assess whether similar effects were observed in human macrophages, THP-1 cells were infected with *B. burgdorferi* in the presence or absence of an IFNAR1-blocking antibody. IFNAR1 blockade significantly reduced IL-6 production at 6- and 24-HPI, and decreased TNF-α levels at early time points (**Fig. 6B**). In contrast, IL-1β production by THP-1 cells was not significantly affected by IFNAR1 blockade. Notably, TNF-α levels in THP-1 cultures were substantially lower than those observed in murine BMDMs, consistent with known species-specific differences in cytokine responses. Overall, these findings indicate that IFN-I signaling modulates pro-inflammatory cytokine production in both murine and human macrophages following *B. burgdorferi* infection.

#### IFN-I signaling promotes pro-inflammatory cytokine expression at the site of *B. burgdorferi* infection *in vivo*

To determine whether IFN-I signaling regulates inflammatory cytokine responses *in vivo* during early *B. burgdorferi* infection, we analyzed cytokine gene expression at the site of infection in WT and *Ifnar1*⁻/⁻ mice. Skin tissues from the inoculation site were collected at 5 DPI. The transcript levels of *Il1b*, *Il6*, and *Tnf* were quantified by quantitative reverse transcription PCR (qRT-PCR) using the primer sequence shown in Table 1. Compared with WT controls, *Ifnar1*⁻/⁻ mice exhibited significantly reduced expression of *Il1b*, *Il6*, and *Tnf* at the site of infection following *B. burgdorferi* challenge (Fig. **7A–C**). These results demonstrate that IFN-I signaling is required for optimal induction of pro-inflammatory cytokine responses *in vivo* during the early stage of infection.

**Fig. 7.**
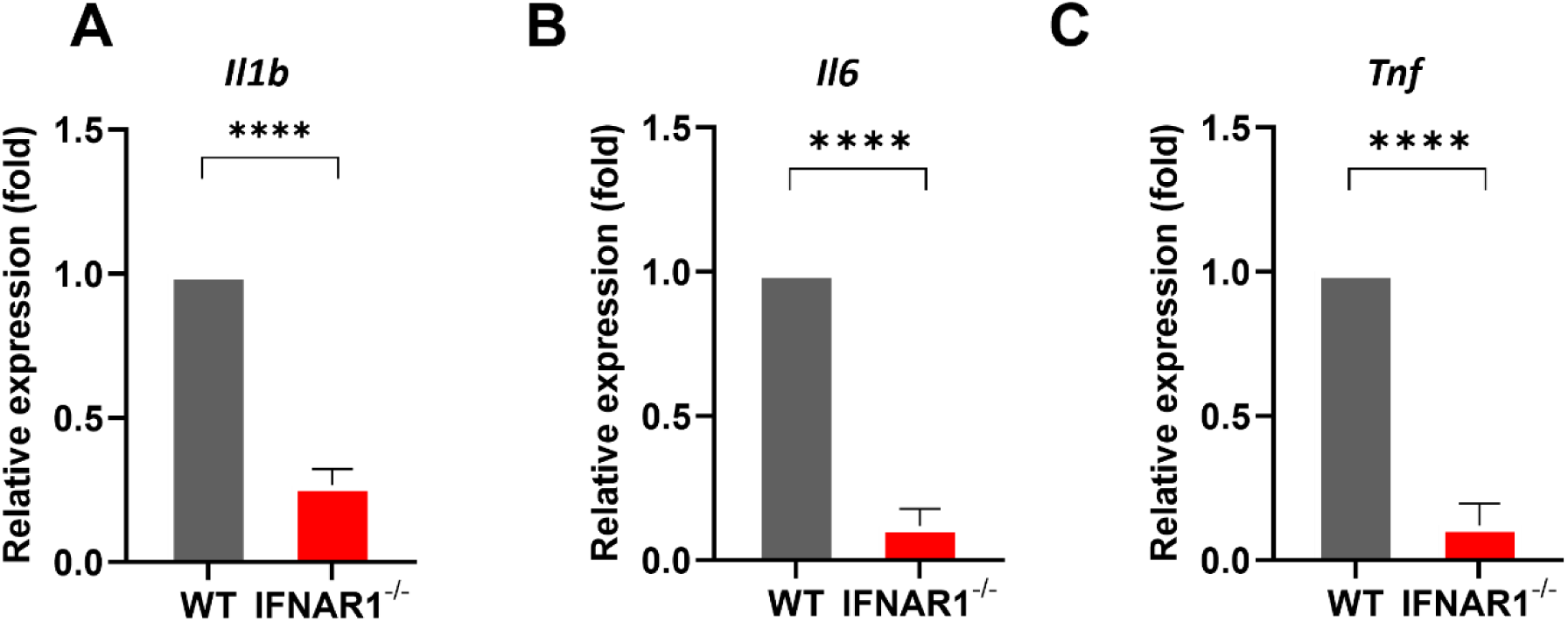
IFN-I signaling promotes pro-inflammatory cytokine expression at the site of *B. burgdorferi* infection. Wild-type (WT) and *Ifnar1*⁻/⁻ mice were infected intradermally with *B. burgdorferi* (1 × 10⁴ spirochetes/ mouse). Skin tissues from the inoculation site were harvested at 5 DPI, and total RNA was isolated. Transcript levels of *Il1b* (**A**), *Il6* (**B**), and *Tnf* (**C**) were quantified by qRT-PCR and normalized to the housekeeping gene *Actin*. Data are presented as mean ± SD from two independent experiments (n = 3–4 mice/ group). Statistical significance was determined using a two-tailed unpaired Student’s *t-*test. ****, *P* < 0.0001.

**Table 1:**
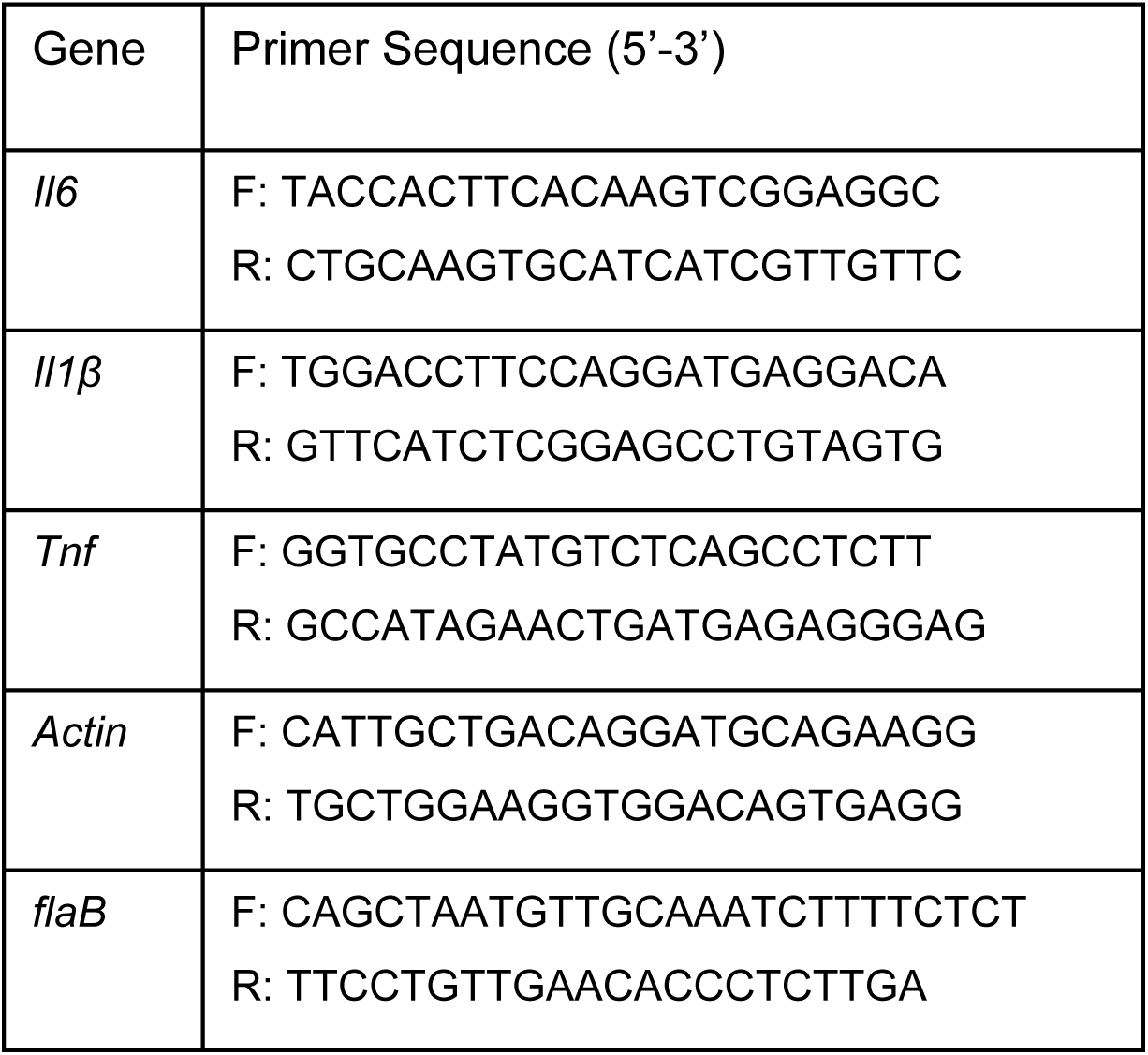
Primer sequence used in this study.

#### Type I IFN signaling promotes recruitment of innate immune cells to the site of *B. burgdorferi* infection

To determine whether IFN-I signaling influences the recruitment of innate immune cells during early *B. burgdorferi* infection, WT and *Ifnar1*⁻/⁻ mice were infected intraperitoneally, and immune cell populations at the site of infection were analyzed 24 HPI by flow cytometry. Using a sequential gating strategy, macrophages were identified as live CD45⁺CD11b⁺F4/80⁺ cells, and neutrophils were identified as live CD45⁺CD11b⁺Ly6G⁺ cells (**Fig. 8A**). Quantitative analysis revealed a significant reduction in the percentages of both macrophages and neutrophils in the peritoneal cavity of *Ifnar1*⁻/⁻ mice compared with WT controls on infection with *B. burgdorferi* (**Fig. 8B**). These results indicate that IFN-I signaling promotes the recruitment of innate immune cells to the site of *B. burgdorferi* infection, thereby contributing to effective early host defense.

**Fig. 8.**
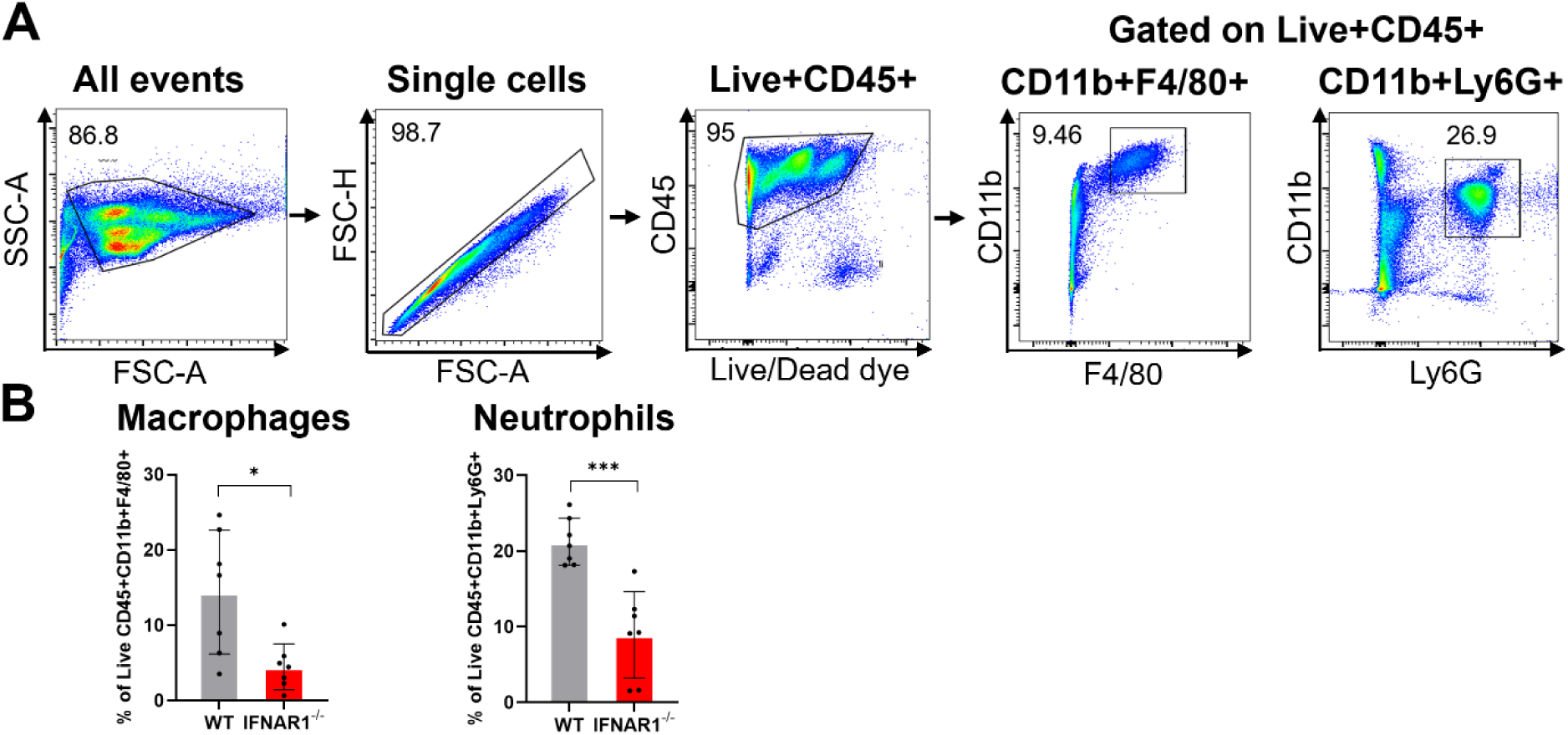
Type I IFN signaling promotes recruitment of innate immune cells during early *B. burgdorferi* infection. Wild-type (WT) and *Ifnar1*⁻/⁻ mice were infected intraperitoneally with *B. burgdorferi* (1 × 10⁴ spirochetes/mouse). Peritoneal exudate cells were harvested at 24 HPI and analyzed by flow cytometry. (**A**) Flow cytometry gating strategy used to identify macrophages (live CD45⁺CD11b⁺F4/80⁺) and neutrophils (live CD45⁺CD11b⁺Ly6G⁺). (**B**) Quantification of the percentages of macrophages and neutrophils in the peritoneal cavity. Data are presented as mean ± SD (n = 6–8 mice/ group) from two independent experiments. Statistical significance was determined using a two-tailed unpaired Student’s *t-*test. *, *P* < 0.05; ***, *P* < 0.001.

### Pharmacological activation of IFN-I signaling enhances *B. burgdorferi* clearance *in vitro* and in vivo

Given that loss of IFN-I signaling impaired phagocytosis, intracellular killing, cytokine production, and immune cell recruitment, we next asked whether pharmacological activation of the IFN-I signaling could enhance host control of *B. burgdorferi*. As bacterial cyclic di-adenosine monophosphate (c-di-AMP) is an activator of cGAS–STING–dependent IFN-I signaling, we used c-di-AMP as a tool to augment IFN-I responses *in vitro* and *in vivo*. Accordingly, RAW 264.7 macrophages were pretreated with c-di-AMP before infection with GFP-expressing *B. burgdorferi* at low and high MOI. The result showed that c-di-AMP pretreatment significantly increased macrophage phagocytosis, as reflected by an increased percentage of GFP-positive cells and elevated GFP MFI compared with untreated controls (**Fig. 9A**). Further gentamicin protection assays demonstrated a reduction in intracellular bacterial viability in c-di-AMP–treated macrophages, indicating enhanced intracellular killing (**Fig. 9B**).

**Figure 9.**
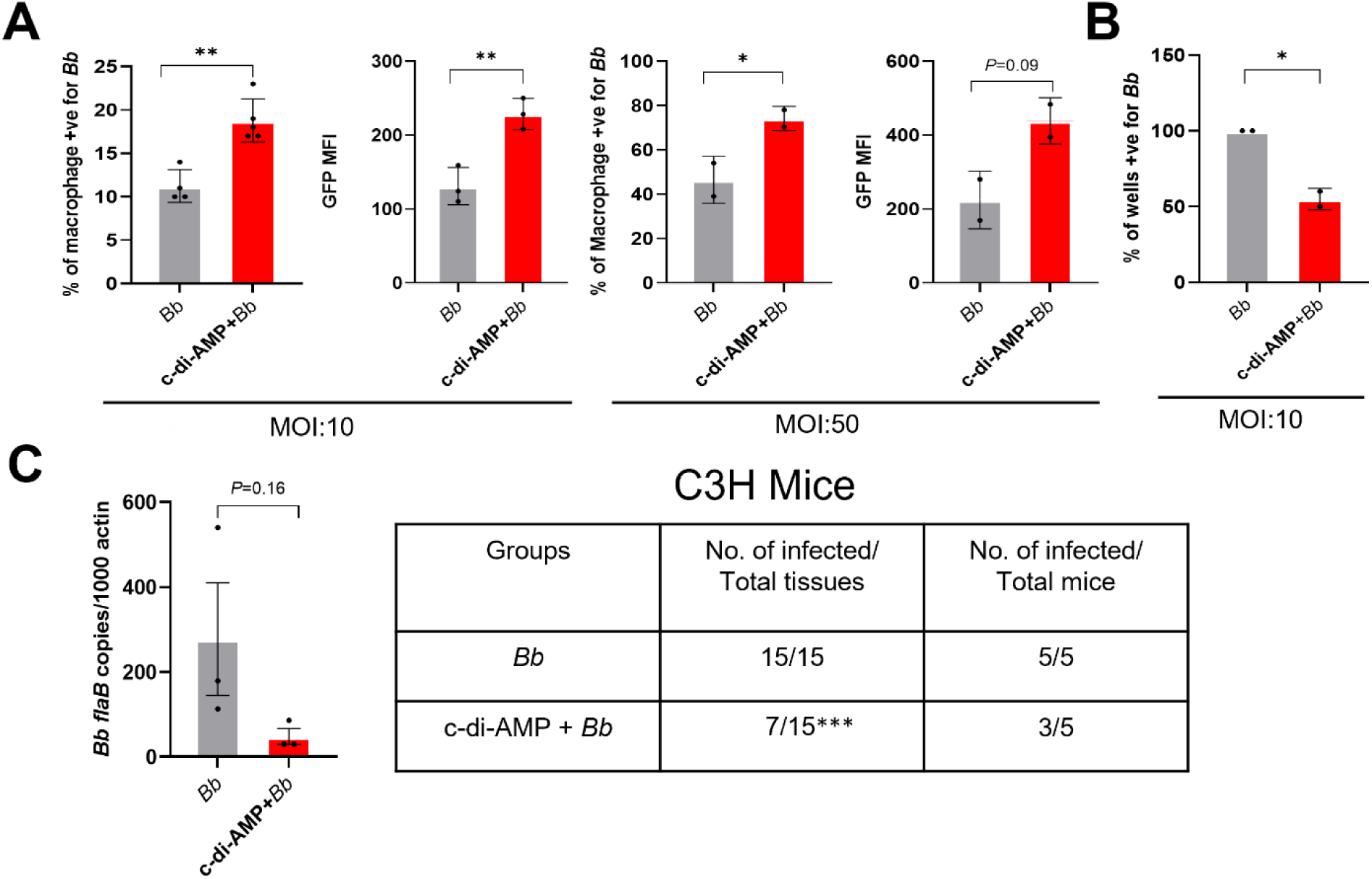
Pharmacological activation of IFN-I signaling enhances *B. burgdorferi* clearance *in vitro* and *in vivo*. **(A)** *In vitro* macrophage assays. RAW 264.7 macrophages were pretreated with c-di-AMP before infection with GFP-expressing *B. burgdorferi* at low and high MOI. Phagocytosis was assessed by flow cytometry to determine the percentage of GFP-positive cells and GFP MFI. (**B**) Gentamicin protection assay. To assess intracellular spirochete viability, RAW 264.7 macrophages were pretreated with c-di-AMP and subsequently infected with *B. burgdorferi* for 2.5 hours, followed by treatment with gentamicin (200 µg/mL) to eliminate extracellular bacteria. Following washing and lysis, cell lysates were diluted in BSK-II medium and plated in 96-well plates. Intracellular bacterial viability was assessed by limiting dilution, and data are presented as percentages of *B. burgdorferi-*positive wells. (**C**) *In vivo* bacterial clearance. Mice were pretreated intradermally with c-di-AMP before infection with *B. burgdorferi* (1Χ10^4^ spirochetes/ mouse). Spirochetal burden at the site of infection was quantified by qPCR at 5 DPI (**left**), and dissemination was assessed by the frequency of culture-positive tissues at 10 DPI (**right**). Data are presented as mean ± SD of two independent experiments (n = 3–5 mice/ group). Statistical significance was determined using a two-tailed unpaired Student’s *t-*test for **(A)** and **(B)** (*, *P* < 0.05; **, *P* < 0.01) and a two-tailed Fisher’s exact test for **(C)** (***, *P* < 0.001).

We next evaluated whether c-di-AMP activation of IFN-I signaling could enhance bacterial control *in vivo*. Mice were pretreated intradermally with c-di-AMP before infection with *B. burgdorferi*. Analysis of skin tissues at the site of infection revealed a reduction in bacterial burden at 5 DPI, as determined by qPCR (**Fig. 9C**, left). Consistent with this finding, tissue culture analyses demonstrated a lower frequency of culture-positive tissues at 10 DPI in c-di-AMP–treated mice compared with controls (**Fig. 9C**, right). Together, these results demonstrate that pharmacological activation of IFN-I signaling enhances phagocytosis and intracellular killing of *B. burgdorferi in vitro* and promotes bacterial clearance *in vivo*.

## DISCUSSION

The prevailing view of Type I interferon (IFN-I) signaling during *B. burgdorferi* infection emphasizes its role in driving pathogenic outcomes, particularly the development and severity of Lyme arthritis, while considering it largely dispensable for host defense [21, 26–29]. This perspective is based primarily on murine studies in which IFNAR1 deficiency reduces joint pathology without significantly affecting spirochetal burdens at four weeks post-infection[29]. In the present study, we uncover a previously underappreciated protective function of IFN-I during the acute phase of infection. By focusing on the earliest window following inoculation, we demonstrate that IFN-I signaling is required to limit spirochetal expansion at the site of infection through enhanced macrophage phagocytosis, intracellular killing, and pro-inflammatory cytokine production. In light of the robust IFN-I signatures documented in human erythema migrans lesions and early disseminated disease [30–32], our findings support a dual, time-dependent role for IFN-I in *B. burgdorferi* infection: an early protective response that restricts bacterial growth, followed by a later contribution to inflammatory joint pathology.

A central mechanistic insight of this work is that IFN-I signaling enhances macrophage-mediated phagocytosis and intracellular killing of *B. burgdorferi*. Phagocytic clearance is a key determinant of infectivity, as the ability of *B. burgdorferi* to establish infection correlates strongly with resistance to elimination by professional phagocytes [35]. Although IFN-I is known to modulate phagocyte function in other bacterial models [10–12, 36, 37], its role in *B. burgdorferi* clearance has not been defined. Phagocytes recognize spirochetes through receptors including TLR2, complement receptor 3 (CR3), and scavenger receptors, activating MyD88-dependent and -independent pathways that drive bacterial uptake and degradation [24, 33, 34, 38–41]. While effective phagocytosis reduces bacterial burden and initiates proinflammatory cytokine production [42], impaired clearance permits *B. burgdorferi* to disseminate and persist [34, 43]. Consistent with this concept, *B. burgdorferi* has evolved mechanisms to evade phagocytic elimination, and its infectivity correlates with resistance to phagocyte-mediated killing [35, 44]. In this study, we show that IFNAR1-deficient macrophages display impaired spirochetal uptake both *in vitro* and *in vivo*, accompanied by defective lysosomal targeting and reduced intracellular killing (**Figs. 2, 3,** and **4**), resulting in a higher *B. burgdorferi* burden at the site of infection and the heart (**Fig. 1**). Although MyD88-dependent signaling contributes to *B. burgdorferi* recognition, it is insufficient for complete bacterial clearance [34, 41, 45]. Our findings, therefore, identify IFN-I signaling as a key MyD88-independent pathway linking intracellular pathogen sensing to effective antimicrobial effector functions. Notably, *B. burgdorferi* linear plasmid lp36 has been shown to be required for both optimal l IFN-I induction and efficient adhesion and internalization, suggesting a functional coupling between IFN-I signaling and phagocytic recognition [46]. In addition, IFN-I signaling supports optimal pro-inflammatory cytokine production (**Figs. 6** and **7**), further amplifying innate immune responses required for early containment of infection.

Beyond macrophage-intrinsic antimicrobial functions, IFN-I also plays a key role in coordinating cellular recruitment to the site of infection. IFNAR1-deficient mice exhibited reduced recruitment of both macrophages and neutrophils during early infection (**Fig. 8**), underscoring the importance of IFN-I in shaping the local inflammatory environment. Early recruitment of these innate immune cells is essential for limiting *B. burgdorferi* replication and dissemination. Previous studies have shown that impaired macrophage or neutrophil recruitment leads to increased spirochetal burden [21, 43, 47–52], whereas enhanced neutrophil chemotaxis markedly attenuates infectivity [53]. In fact, tick saliva actively compromises neutrophil function to facilitate initial infection and dissemination [54, 55]. Together, these observations position IFN-I as an upstream regulator that integrates innate cell recruitment with antimicrobial function to ensure early bacterial containment.

The protective role of IFN-I in early spirochetal control uncovered in this study gains additional significance in the context of natural tick-borne transmission, where successful infection depends critically on immune evasion. *B. burgdorferi* employs a dual-pronged suppression strategy, leveraging both tick saliva components, such as the salivary protein Salp15, which inhibits immune cell activation, and spirochetal virulence factors that dampen host IFN-I signaling in the skin [56–58]. The protective nature of the host IFN-I response is most clearly illustrated by studies of the *B. burgdorferi* virulence factor BBA57. Wild-type spirochetes use BBA57 to suppress expression of the host IFN-I gene *Ifna9* (interferon-α9). In contrast, *bba57* mutant spirochetes fail to suppress *Ifna9*, leading to enhanced IFN-I activation, impaired dissemination, and efficient clearance at the inoculation site [58]. This genetic evidence demonstrates that IFN-I activation is inherently protective and must be actively suppressed by the pathogen to establish a productive, disseminating infection. Our findings reinforce this paradigm by showing that, when the tick-mediated immune-evasive environment is bypassed, IFN-I signaling is indispensable for effective early clearance of *B. burgdorferi*.

The ability of exogenous IFN-I to enhance macrophage antimicrobial capacity represents an important and potentially translational aspect of our findings. Short-term pretreatment with IFN-β significantly increased both uptake and intracellular degradation of *B. burgdorferi* (**Fig. 5**), indicating that IFN-I can directly prime phagocytes for enhanced antimicrobial function. This priming effect is likely mediated through the induction of interferon-stimulated genes (ISGs) that potentiate cytoskeletal remodeling, phagosome maturation, and lysosomal killing. Extending this concept, we demonstrate that pharmacological activation of IFN-I signaling using the bacterial second messenger c-di-AMP markedly enhances phagocytosis and intracellular killing *in vitro* and restricts bacterial survival *in vivo* (**Fig. 9**). These results help reconcile prior reports describing neutral or detrimental effects of IFN-I in Lyme disease by highlighting the critical importance of timing and context [21, 45]. Early IFN-I signaling enhances antimicrobial defenses, whereas prolonged or dysregulated IFN-I responses later in infection may suppress protective immunity and promote inflammatory pathology.

Several limitations of this study should be acknowledged. Our analyses focused primarily on macrophages and early innate responses; whether IFN-I similarly regulates antimicrobial functions in other phagocytes, such as dendritic cells, remains to be determined. In addition, our use of needle inoculation does not fully recapitulate the complex immunomodulatory environment of natural tick-borne transmission. Future studies employing tick-mediated infection models will be important to validate and extend these findings. Finally, the long-term consequences of modulating IFN-I signaling on bacterial persistence and chronic inflammatory outcomes warrant further investigation.

In summary, our findings support a model in which IFN-I signaling exerts temporally distinct effects during *B. burgdorferi* infection (**Fig. 10**). During the earliest phase of infection, IFN-I acts as a pivotal amplifier of innate immunity, enhancing phagocytosis, promoting intracellular killing, augmenting pro-inflammatory cytokine production, and facilitating recruitment of macrophages and neutrophils, thereby constraining bacterial expansion and early dissemination. In contrast, sustained or excessive IFN-I signaling at later stages contributes to inflammatory pathology, including Lyme arthritis [21]. This temporally integrated framework reconciles previously perceived pathogenic roles of IFN-I with its essential early protective function. Collectively, these findings suggest that precisely timed modulation of IFN-I pathways may enhance early host defense while limiting downstream immunopathology.

**Figure 10.**
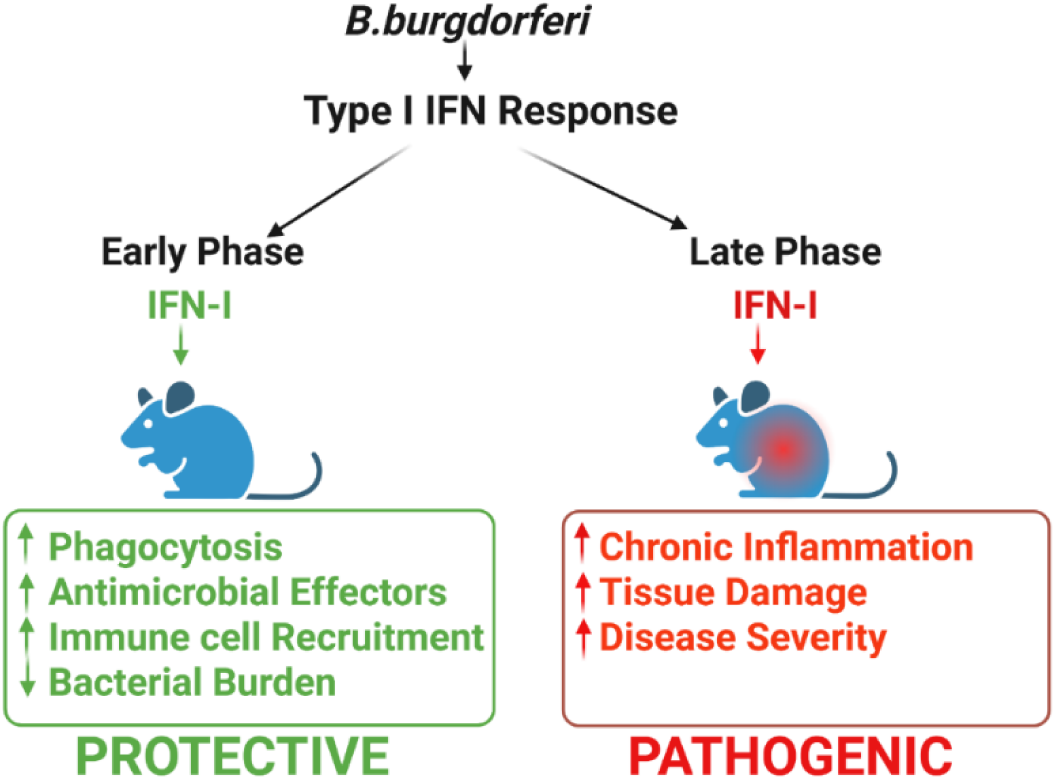
Proposed model illustrating dual roles of Type I IFN signaling during *B. burgdorferi* infection. Early during infection, *B. burgdorferi* induces IFN-I signaling, which promotes protective host defense by enhancing macrophage phagocytosis, facilitating intracellular killing through lysosomal targeting, inducing pro-inflammatory cytokine production, and supporting recruitment of innate immune cells to the site of infection. These coordinated responses control spirochetal loads at the inoculation site. During later stages of infection, sustained or dysregulated IFN-I signaling contributes to inflammatory pathology, including the development of Lyme arthritis. Together, this model highlights the context- and time-dependent nature of IFN-I signaling during *B. burgdorferi* infection, in which early IFN-I responses are protective, whereas prolonged IFN-I activity contributes to disease pathology.

## MATERIALS AND METHODS

### Ethics statement

All animal experiments were approved by the Indiana University School of Medicine’s Institutional Animal Care and Use Committee and were performed following the institutional guidelines for animal use. Mice were housed at the Laboratory Animal Research Centre (LARC) facility.

### Mice

Wild-type (WT) C3H/HeN, C57BL/6, and *Ifnar1*⁻/⁻ mice (on a C57BL/6 background), were purchased from the Jackson Laboratory (Bar Harbor, ME) and bred in-house at the Indiana University LARC facility. Both male and female mice aged 4 to 8 weeks were used for all the experiments. Mice were housed under specific pathogen-free conditions in filter-top cages with a 12-hour light/dark cycle and provided food and water *ad libitum*.

### *B. burgdorferi* strains and culture conditions

The virulent *B. burgdorferi* strain 5A4NP1 was used for all the infection studies. A GFP-expressing strain was generated by electroporation of the pTM61-GFP plasmid, which drives GFP expression under the control of the *flaB* promoter. *B. burgdorferi* strains were cultured in complete Barbour-Stoenner-Kelly II (BSK-II) medium supplemented with 6 % normal rabbit serum at 37°C with appropriate antibiotics. Antibiotics were used at the following concentrations: gentamicin at 50 µg/mL, and kanamycin at 100 µg/mL. Cultures were visually monitored for spirochete motility and density by dark-field microscopy.

### Cell lines

Murine macrophage cells RAW 264.7 (ATCC, Manassas, VA) were cultured in Dulbecco modified Eagle medium (DMEM) with L-glutamine supplemented with 10% heat-inactivated FBS and 1% penicillin-streptomycin. THP-1 human monocyte cells were used from our lab inventory and cultured in RPMI 1640 containing 10% FBS. THP-1 cells were differentiated into macrophage-like cells using phorbol 12-myristate 13-acetate (PMA;10 ng/mL) for 24 hours prior to infection. L929 mouse fibroblasts were maintained in DMEM supplemented with 10% FBS using standard culture conditions.

### Preparation of Bone Marrow-Derived Macrophages (BMDMs)

Primary bone marrow-derived macrophages (BMDMs) were generated from femurs of 4-6-week-old WT and *Ifnar1*⁻/⁻ mice as described earlier [59]. In brief, bone marrow cells were flushed from mouse femurs with sterile RPMI medium and cultured for 7 days in RPMI supplemented with 30% L929 cell-conditioned medium, 20% fetal bovine serum (FBS), and 1% penicillin-streptomycin. Differentiated macrophages were harvested and seeded into tissue culture plates for subsequent experiments.

### Mouse infection and *B. burgdorferi* burden determination by qPCR

WT and *Ifnar1*⁻/⁻ mice were infected intradermally with *B. burgdorferi* (1Χ10^4^ spirochetes/ mouse). At 5- and 10-DPI, mice were euthanized, and skin (site of infection), heart, and joint tissues were harvested. Genomic DNA was isolated using the DNeasy Blood & Tissue Kit (Qiagen) according to the manufacturer’s instructions. *B. burgdorferi* burden was quantified by quantitative PCR (qPCR) targeting the *B. burgdorferi flaB* gene, with results normalized to mouse *β-actin*. Results are expressed as *flaB* copies per 10^6^ *β-actin* copies. Primer sequences are listed in Table 1.

To assess bacterial viability, tissues were cultivated in BSKL-II medium supplemented with phosphomycin (2 mg/mL), rifampin (5 mg/mL), and amphotericin B (250 µg/mL) (Sigma-Aldrich). All cultures were incubated at 37°C and monitored for spirochete growth by dark-field microscopy for up to 21 days. A tissue was scored as positive if at least one motile spirochete was detected during the observation period.

### *In vitro* Phagocytosis Assay

Macrophages (1Χ10^6^ cells/well, 12-well plates) were infected with GFP-expressing *B. burgdorferi* at an MOI of 10. Where indicated, cells were pretreated for 1 hour with recombinant IFN-β (500 pg/mL, Sino Biological Inc., 50708) or an anti-mouse IFNAR1 blocking antibody (1µg/mL, BioLegend, 127322, clone MAR1-5A3), with isotype-matched IgG as a control (BioLegend, 400166, clone MOPC-21). Infections were synchronized at 4°C for 15 minutes, followed by incubation at 37°C for 2.5 hours. Cells were washed extensively with cold PBS to remove extracellular bacteria, resuspended in FACS buffer (PBS containing 2% FBS), and analyzed by flow cytometry (Fortessa, BD Bioscience). Mock-infected cells and cells infected with unlabeled *B. burgdorferi* served as negative controls to set the GFP negative gate. Macrophages were gated based on forward (FSC) and side scatter (SSC) to exclude cellular debris and doublets. Data was analyzed using FlowJo software (version 10, BD Life Sciences). Phagocytosis was quantified as the percentage of GFP-positive cells and GFP mean fluorescence intensity (MFI).

### *In vivo* Phagocytosis Assay

WT and *Ifnar1*⁻/⁻ mice were infected intraperitoneally (IP) with GFP-expressing *B. burgdorferi* (1×10⁷ spirochetes/ mouse) or non-fluorescent *B. burgdorferi* (used to set the GFP negative gate and control for autofluorescence) for 6 hours as described previously [34]. Peritoneal exudate cells (PECs) were collected and stained with LIVE/DEAD dye (Fixable viability Dye eFluor 780, Invitrogen), followed by blocking with CD16/32 (BD Pharmingen, 553142) for 15 minutes at 4°C. The cells were then stained with anti-CD45 (clone: 30-F11,103133), anti-CD11b (clone: M1/70,101215), and anti-F4/80 (clone: BM8, 123109) antibodies (Abs) for 30 minutes at 4°C in the dark. After staining, cells were washed and fixed in 1% paraformaldehyde for 15 minutes at RT. Fixed cells were resuspended in FACS buffer and subjected to flow cytometry analysis (Fortessa, BD Biosciences). Phagocytosis was assessed by determining the percentage of live CD45^+^CD11b^+^F4/80^+^ cells positive for GFP signal. Single-stained, FMO (Fluorescence minus one), and unstained controls were used to set gating parameters and compensation. Data were analyzed using FlowJo software (version 10, BD Life Sciences).

### Confocal microscopy

Macrophages were seeded at 2 × 10^5^ cells/well on glass coverslips in 12-well cell culture plates (Sigma-Aldrich) and incubated overnight. The following day, cells were washed and infected with GFP-*B. burgdorferi* at an MOI of 10 for 2.5 hours. For some experiments, cells were pre-treated with either recombinant IFN-β (500 pg/mL) or an IFNAR1 blocking antibody (1µg/mL) for 1 hour at 37°C prior to infection. Following extensive washing with ice-cold PBS, cells were fixed in 4% paraformaldehyde. For cytoskeletal staining, cells were permeabilized with 0.1% Triton X-100 and incubated with Phalloidin-California Red Conjugate (1:1000 dilution; Cayman Chemical, 20546) for 1 hour. To assess the colocalization of internalized spirochetes with lysosomes, fixed and permeabilized cells were incubated with an anti-LAMP1 antibody (Cell Signaling Technology, 99437) overnight at 4°C. Subsequently, cells were washed and incubated with an anti-rabbit Alexa Fluor 594-conjugated secondary antibody (Cell Signaling Technology, 35363) for 2 hours at room temperature in the dark. Coverslips were mounted on glass slides using Antifade Mounting Medium containing DAPI (Vector Laboratories, H-1200). Confocal imaging was performed using a Leica upright confocal microscope and LAS X software. Images were acquired using a 40x oil-immersion objective. All imaging was conducted at the Indiana University Microscopy Core Facility.

### Gentamicin protection assay

To assess intracellular killing, a gentamicin protection assay was performed as previously described, with some modifications [60]. Briefly, macrophages were infected with *B. burgdorferi* for 2.5 hours, after which the culture supernatant was collected. Cells were washed gently with ice-cold PBS and resuspended in fresh DMEM containing 10% FBS. Gentamicin (200 µg/mL) was added to the collected supernatant and the infected macrophages, followed by incubation for 4 hours at 37°C to eliminate extracellular bacteria. To confirm the efficiency of the antibiotic treatment, 100 µL of the treated supernatant was diluted in 20 mL of BSK-II medium and plated in 96-well plate. To quantify viable intracellular bacteria, the infected macrophages were washed, incubated with sterile pre-warmed distilled water (200 µL) for 5 minutes, and gently lysed via pipetting to release internalized *B. burgdorferi*. A 100 µL of the resulting cell lysate was resuspended in 20 mL of BSK-II medium containing phenol red and kanamycin and plated in 96-well plates. The plates were incubated at 37°C for 21 days. Final growth determination was based on the color change of the medium and confirmed by the presence of motile spirochetes under dark-field microscopy.

### Cytokine analysis by ELISA

To assess cytokine production, BMDMs from wild-type (WT) and *Ifnar1*⁻/⁻ mice, as well as THP-1 cells were infected with live *B. burgdorferi* at an MOI of 10 for the indicated time points (3, 6, and 24) hours. For human macrophage experiments, THP-1 cells were pre-treated with either an anti -hIFNAR1 antibody (1µg/mL, InvivoGen, hifnar-mab1) or an isotype control (β-Gal-hIgG1; InvivoGen, bgal-mab1) for 1 hour before infection. Following incubation, cell supernatants were collected and centrifuged to remove cellular debris. Concentrations of TNF-α, IL-6, and IL-1β in the supernatant were quantified using commercial ELISA kits (BioLegend; murine:430904, 431304, 432604; human: 430204, 430504, 437004) according to the manufacturer’s instructions.

### Quantitative RT-PCR Analysis

Total RNA was isolated from mouse skin tissues using TRIzol reagent (Thermo Fisher Scientific) according to the manufacturer’s protocol. To eliminate genomic DNA (gDNA) contamination, all samples were subjected to on-column digestion using the RNase-free DNase set (Qiagen, 79254), and the RNA was subsequently purified using the RNeasy Mini Kit (Qiagen, 74106). RNA purity and concentration were determined using a NanoDrop One spectrophotometer (Thermo Fisher Scientific). cDNA was synthesized from the RNA template using the PrimeScript 1st strand cDNA Synthesis Kit (Takara Bio, 6110A) according to the manufacturer’s instructions. Quantitative PCR (qPCR) was performed in triplicate technical replicates using PowerUp SYBR Green Master Mix (Thermo Fisher Scientific, A25742) on an ABI 7000 Sequence Detection system (Applied Biosystems). The specific primer sequences are detailed in Table 1. The mRNA expression levels of target genes were normalized to the expression of the housekeeping gene *β-actin*. Relative gene expression was calculated using the comparative ΔΔC_T_ method (2^-ΔΔCT^) as described previously [61].

### Immune cell recruitment analysis

WT and *Ifnar1*⁻/⁻ mice were infected intraperitoneally with *B. burgdorferi* (1 × 10^4^ spirochetes/mouse). At 24 HPI, mice were euthanized, and peritoneal exudate cells (PECs) were harvested. Total cell counts were determined using Trypan blue exclusion. For flow cytometric analysis, PECs were stained with a fixable viability dye (eFluor 780; Invitrogen), followed by incubation with CD16/32 blocking antibody (clone 2.4G2; BD Pharmingen, 553142) for 15 minutes at 4°C. Cells were then stained with anti-CD45 (clone 30-F11; BioLegend, 103133), anti-CD11b (clone M1/70; BioLegend, 101215), anti-F4/80 (clone BM8; BioLegend, 123109), and anti-Ly6G (clone 1A8; BioLegend, 127613) antibodies for 30 minutes at 4°C in the dark. Following staining, cells were washed with PBS and fixed in 1% paraformaldehyde for 15 minutes at room temperature. Fixed cells were resuspended in FACS buffer and subjected to flow cytometry analysis (Fortessa, BD Biosciences). Single-stained, FMO, and unstained controls were used to set gating parameters and compensation. Data were analyzed using FlowJo software (version 10, BD Life Sciences).

### Agonist treatment analysis

For *in vitro* experiments, RAW 264.7 macrophages were pre-treated with c-di-AMP (1µg/mL; InvivoGen, tlrl-nacda) for 1 hour before infection with GFP-expressing or non-fluorescent *B. burgdorferi*. The percentage of GFP-positive macrophages and the GFP MFI were determined by flow cytometry. Intracellular bacterial viability was assessed via a gentamicin protection assay as described above.

To elucidate the effect of c-di-AMP *in vivo*, mice were pre-treated intradermally with c-di-AMP (50 µg/mouse; InvivoGen, tlrl-nacda) 24-hours before infection with *B. burgdorferi* (1 × 10^4^ spirochetes/mouse). *B. burgdorferi* burden at the site of infection was determined by qPCR, and the presence of viable bacteria was determined by culturing tissues in BSK-II BAM medium.

### Statistical Analysis

Statistical analyses were performed using Prism (GraphPad Software). All the experiments were performed at least twice. Data are presented as mean ± standard deviation (SD). For comparisons between two groups, statistical significance was evaluated using an unpaired, two-tailed Student’s t-test. For categorical data, statistical comparisons were performed using a two-tailed Fisher’s exact test. *P*-values are represented as \**P*<0.05; \*\**P*<0.01; \*\*\**P*<0.001; \*\*\*\**P*<0.0001; ns, not significant.

## ACKNOWLEDGMENTS

This research was partly funded by NIH grants R01AI152235 (to X. F. Yang, H. O. Sintim) and R01AI083640 (to X. F. Yang).

## Abbreviations

Bb: Borrelia burgdorferi
IFN-I: Type I Interferon
STING: Stimulator of Interferon Genes
MFI: Mean Fluorescence Intensity
DPI: Days Post-infection
HPI: Hours Post-infection

